# Glucose modulates *Drosophila* longevity and immunity independent of the microbiota

**DOI:** 10.1101/027326

**Authors:** Anthony Galenza, Jaclyn Hutchinson, Bart Hazes, Shelagh D. Campbell, Edan Foley

**Affiliations:** University of Alberta, Department of Medical Microbiology and Immunology, Institute of Virology.

**Keywords:** *Drosophila*, nutrition, microbiome, protein to carbohydrate, ageing

## Abstract

The acquisition of nutrients is essential for maintenance of metabolic processes in all organisms. Nutritional imbalance contributes to myriad metabolic disorders that include malnutrition, diabetes, and even cancer. Recently, the importance of macronutrient ratio of food has emerged as a critical factor to determine health outcomes. Here we show that individual modifications to a completely defined diet markedly impact multiple aspects of organism wellbeing in *Drosophila melanogaster.* Through a longitudinal survey of several diets we demonstrate that increased levels of dietary glucose significantly improve longevity and immunity in adult *Drosophila.* Our metagenomic studies, show that relative macronutrient levels not only influence the host, but also have a profound impact on microbiota composition. However, we found that elevated dietary glucose extended the lifespan of adult flies even when raised in a germ-free environment. Furthermore, when challenged with a chronic enteric infection, flies fed a diet with added glucose had increased survival times even in the absence of an intact microbiota. Thus, in contrast to known links between the microbiota and animal health, our findings uncover a novel microbiota-independent response to diet that impacts host wellbeing. As dietary responses are highly conserved in animals, we believe our results offer a general understanding of the association between glucose metabolism and animal health.

## INTRODUCTION

Recent developments in the production, distribution and consumption of food fundamentally transformed our relationship with our nutritional environment. The near limitless availability of ready-made, high-calorie meals is a prominent contributor to the emergence of metabolic disorders as a major health challenge in many nations, and there is an increased emphasis on the importance of nutritional awareness to optimize individual health outcomes (Simpson et al., 2015). In this context, the obvious health benefits of a nutritionally replete diet fuel a multi-billion dollar health and nutrition industry that centers on the pursuit of a “balanced” diet. However, functional definitions of health and nutritional balance are more complex than may seem apparent and require more than a steady intake of specific amounts of nutrients.

Dietary influence on longevity has been extensively studied in several vertebrate and invertebrate models (Fontana and Partridge, 2015; Tatar et al., 2014). Initial, widely reported observations showed that caloric restriction promotes a longer lifespan in rats, and this was supported by subsequent studies in mice, *Drosophila*, worms, and yeast (Dilova et al., 2007; Fontana et al., 2010). Recent long-term experiments yielded mixed observations on the benefits of caloric restriction for primates (Colman et al., 2014; Mattison et al., 2012). However, the two studies in question differed considerably in their experimental protocols, making comparisons difficult. Exploration of the basis for extended longevity in models of caloric restriction emphasized the relative contributions of individual nutrients to animal lifespan (Lee et al., 2015; Simpson and Raubenheimer, 2009; Solon-Biet et al., 2015b). These studies revealed that diets with low protein to carbohydrate ratios significantly extended the lifespans of mice and *Drosophila* (Lee et al., 2008; Solon-Biet et al., 2014). More recent studies have focused on additional dietary contributions to health and lifespan that include the time of consumption (Jakubowicz et al., 2013; Mattson et al., 2014), duration of fasting between meals (Harvie et al., 2011; Honjoh et al., 2009), as well as relative amounts of amino acids in the diet (Miller et al., 2005; Wu et al., 2013). When considered as a whole, these studies point to a remarkably nuanced relationship between the uptake of dietary nutrients and animal wellbeing.

Many studies of the interplay between nutrition and health overlook microbial contributions. In particular, we know very little about the relationship between the intestinal microflora, host diet, and host intestinal immunity. We consider this a particularly relevant aspect of health and lifespan, as diet and health are intimately linked by the intestinal microbiota (Flint et al., 2012). Diet shapes the composition of the intestinal microflora, which, in turn, influences events as diverse as nutrient allocation, intestinal physiology, immune responses, and the onset of chronic diseases. For example, the intestinal microbiota facilitates the uniquely restricted diet of the koala (Osawa et al., 1992); orchestrates the establishment of immune structures in mammals (Hooper et al., 2012); and contributes to the containment, or dissemination of pathogenic microbes in a number of experimental models (Round and Mazmanian, 2009; Wlodarska et al., 2015). The genetically accessible model system *Drosophila melanogaster* is a particularly valuable tool to reveal key aspects of relationships between diet, the microbiota and the host (Erkosar and Leulier, 2014; Ma et al., 2015). The fly gut shares numerous similarities with mammalian counterparts that include developmental origin, cellular composition, and metabolic pathways (Lemaitre and Miguel-Aliaga, 2013). Additionally, while the mammalian gut contains 500-1000 separate bacterial species, the fly gut is far simpler to study with 5-30 aerotolerant, cultivable commensal species (Broderick and Lemaitre, 2012; Buchon et al., 2013). A number of recent publications established clear mechanistic relationships between the intestinal microflora of flies and events as diverse as nutritional regulation (Newell and Douglas, 2014; Storelli et al., 2011; Wong et al., 2014), activation of progrowth pathways (Shin et al., 2011; Storelli et al., 2011), control of immune pathways (Broderick et al., 2014; Erkosar et al., 2014), and selection of mates (Sharon et al., 2010).

Previous studies with *Drosophila* as a tool to explore host-diet-microbiota relationships relied on partially defined oligdic diets. Recently, Piper *et al.* established a protocol to prepare a holidic diet for *Drosophila*, in which the exact composition and concentration of every ingredient is known (Piper et al., 2014). This allows for precise manipulation of nutrient availability in dietary studies, as individual components can be modified to a specified quantity and effects on the organism can be observed.

In this study, we investigated how dietary modifications, inspired in part by popular human diets, affect the health of a fly. Specifically, we made five separate modifications to the original holidic recipe that include the addition of supplementary glucose, starch, casein, palmitic acid, or ethanol. Respectively, these additions represent diets with higher levels of simple sugar, complex sugar, protein, saturated fatty acids, or alcohol. We investigated several aspects of overall health and nutrition and found that relatively modest dietary modifications exert profound impacts on the lifespan, immune response, and microfloral composition of the host. Of the five dietary modifications tested, we found that the elevation of dietary glucose emerged as the most beneficial manipulation, with effects that included an extended lifespan, increased locomotion, and enhanced immunity against an enteric pathogen. We were particularly intrigued by the relationship between diet, the microbiome, longevity and immunity, as this issue has not been tackled in a systematic study to date. We found that dietary supplementation of glucose greatly increased the diversity of the intestinal microbiome. However, when we eliminated the microbiota from flies, we found that the health benefits of increased glucose were largely independent of the microbiota. Combined, our observations establish that elevated levels of dietary glucose provide numerous benefits to fly health and immunity, and that these benefits do not require an intestinal microbiota.

## RESULTS

### Diet and Age Modify Adult Metabolism in *Drosophila.*

We initially measured the relationship between age, diet and metabolism in adult flies. For these assays, we raised flies on a recently described holidic diet, or a holidic diet supplemented with glucose, starch, casein, palmitic acid, or ethanol. The supplementary regimes allowed us to interrogate the impacts of increased levels of simple or complex sugars, protein, saturated fatty acids, or moderate amounts of alcohol on a common experimental model. We measured the weight, protein content, triglyceride levels and glucose levels of male and female flies raised on the respective diets for five days, ten days, or twenty days. We found that age exerted a significant influence on the weight, protein content, and triglyceride content of adult flies, while diet exerted moderate effects on protein levels (Figure 1A and B). In contrast, we found that diet significantly affected triglyceride and glucose levels in flies. Specifically, we found that supplementation of a holidic diet with extra glucose greatly increased triglyceride and glucose levels in older flies compared to age-matched controls raised on the holidic diet (Figure 1C). These data suggest that increased availability of dietary glucose elevates energy stores, particularly in older flies, without significant effects on weight or protein content.

**Figure 1:**
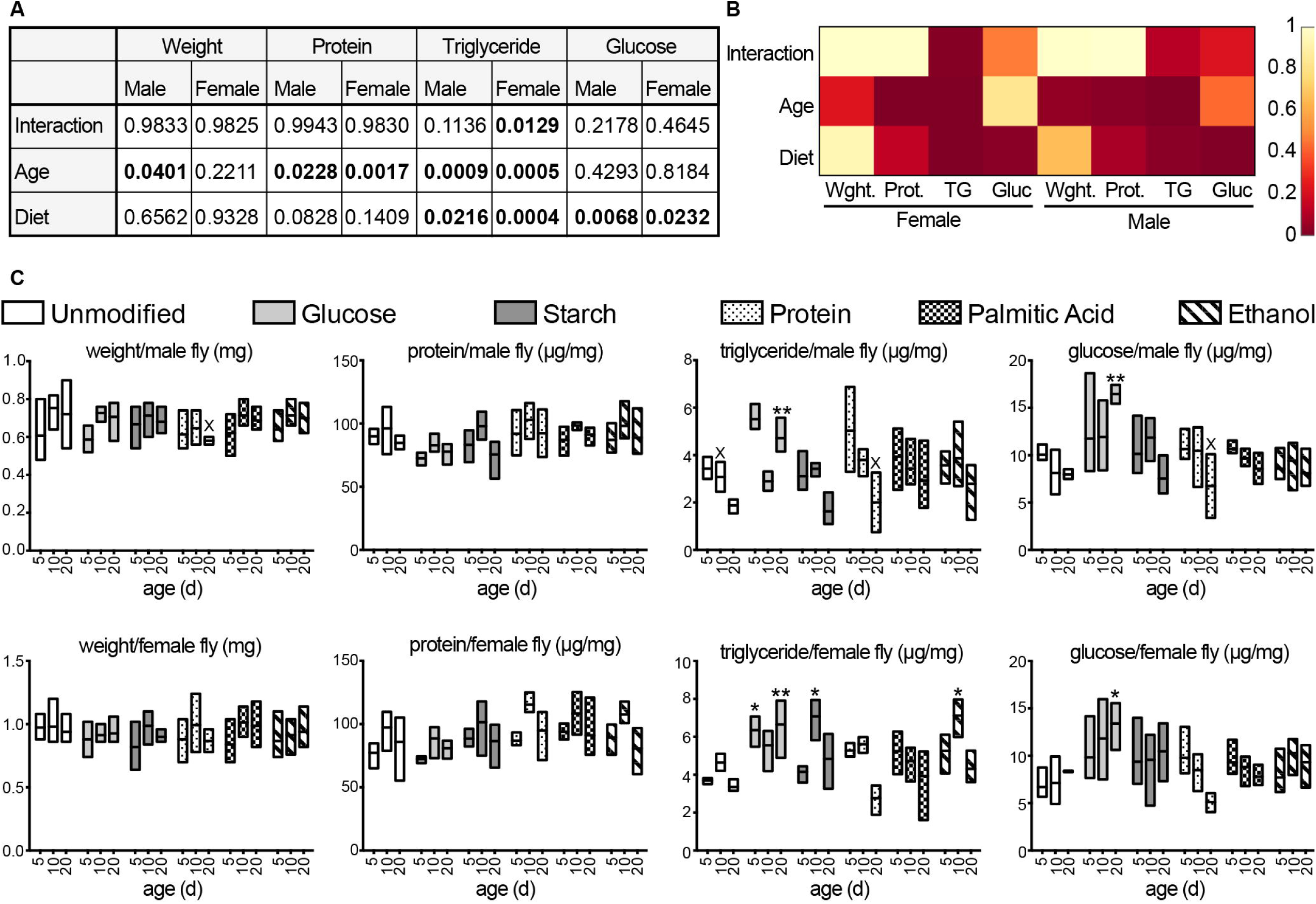
(A) Two-way ANOVA analysis of data from C. Significant P values are highlighted in bold typeface. (B) Heat map summary of P values from A. (C) Longitudinal analysis of weight, protein, triglyceride and glucose content in male and female flies fed the indicated diets. Each column shows the result of three separate measurements at the indicated times, except for columns indicated with an “X”, which show the values of two separate measurements. Mean values for each diet and time point were compared with the means of unmodified diets at the same time with a Bonferroni correction for multiple comparisons. *, p < 0.05; ** p < 0.01.

### Elevated Glucose Availability Extends Adult Longevity

Our data overlap with previous suggestions that dietary modifications have considerable impacts on the metabolic profile of flies (Wong et al., 2014). Numerous studies implicate the availability of nutrients and calories in the control of animal longevity, with a frequent implication that caloric or dietary restriction extends life. However, recent studies also suggest that relative amounts of macronutrients in the diet are important determinants of *Drosophila* lifespan (Lee et al., 2008). Importantly, this hypothesis has not been tested with a defined diet in *Drosophila.* To address this issue, we determined the lifespans of adult male and female flies raised under defined dietary conditions. We found that dietary modifications had slightly different effects on the longevity of male and female flies (Table 1, Figure 2A). In general, dietary modifications that diminished lifespans, such as supplementation with palmitic acid or protein, had more pronounced effects on female flies than male flies, while dietary modifications that extended lifespans, such as addition of ethanol or glucose had more pronounced effects on male flies than females (Figure 2A). We found that elevated glucose availability had a particularly marked impact on longevity in male flies, with a median lifespan extension of 31%. A recent metaanalysis suggested that the longevity benefits of dietary restriction are adaptations to laboratory culture, not a physiological response observed in the wild (Nakagawa et al., 2012). To test if the benefits of glucose addition are restricted to lab-raised flies, we fed adult male *Drosophila melanogaster* captured in the wilds of Edmonton (Canada) an unmodified diet or one supplemented with glucose. As with our lab strains, we found that elevated levels of dietary glucose significantly increased the lifespan of wild flies (Figure 2B and D).

**Table 1:** Median survivals of male and female adult *Drosophila* raised on the respective diets. Each table corresponds to a group of experiments that were performed at the same time and under identical experimental conditions. For each sub-table, Chi square, and P values were calculated with GraphPad Prism 6.0 and are reported relative to the corresponding control flies raised on the holidic diet.

| Diet | Median survival | Chi square | P value |
| --- | --- | --- | --- |
| Holidic | 26 |  |  |
| Starch | 29 | 0.0005998 | 0.9805 |
| Protein | 25 | 2.279 | 0.1312 |
| Palmitic acid | 25 | 0.8032 | 0.3701 |
| Holidic | 26 |  |  |
| Ethanol | 28 | 3.908 | 0.0480 |
| Glucose | 34 | 34.35 | <0.0001 |

FEMALES
| Diet | Median survival | Chi square | P value |
| --- | --- | --- | --- |
| Holidic | 38 |  |  |
| Starch | 35.5 | 3.685 | 0.0549 |
| Protein | 35 | 6.332 | 0.0119 |
| Palmitic acid | 33 | 8.796 | 0.0030 |
| Holidic | 31 |  |  |
| Ethanol | 31.5 | 0.8556 | 0.3550 |
| Glucose | 33 | 6.832 | 0.0090 |

**Figure 2:**
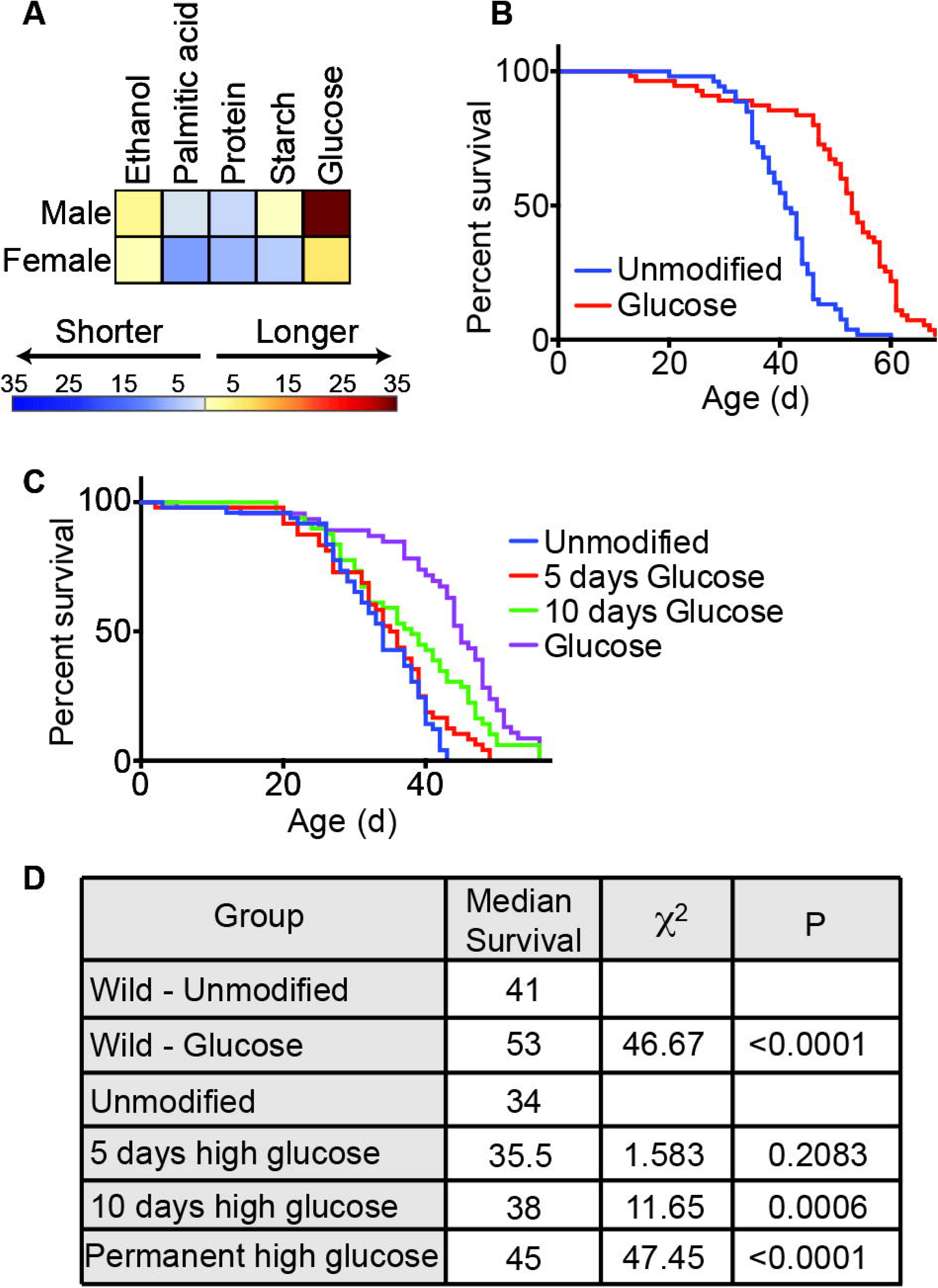
(A) Heat map summary of Chi-square values from table 1 showing the effects of the respective diets on the longevity of male and female flies. Longer values correspond to extended lifespans, and shorter values correspond to diminished lifespans. (B) Survival curves of a wild strain of male *Drosophila* raised on an unmodified diet or on a diet supplemented with glucose. (C) Survival curves of male *Drosophila* raised on an unmodified diet, or on a diet supplemented with glucose for 5 days, 10 days, or permanently. (D) Results of Log-rank (Mantel-Cox) test of data in panel B and C. All □^2^ and p values are relative to unmodified.

Restoration of a complete diet reverts lifespan-extension benefits of dietary restriction In *Drosophila* (Mair et al., 2003). To determine if the benefits of glucose were permanent or transient, we measured the longevity of male flies raised on a holidic diet, or male flies raised on a holidic diet supplemented with extra glucose for the first five days, the first ten days, or the duration of adult life. Our results show that longer periods of dietary supplementation with glucose have more significant effects on lifespan (Figure 2C and D). For example, supplementation of the adult diet with glucose for the first ten days of life extended median survival rates by 12%, while permanent addition of extra glucose extended median survival rates by 32%. These data suggest that overall levels of dietary glucose reversibly influence the lifespan of adult *Drosophila.*

### Elevated Levels of Dietary Glucose Promote Immunity Against an Intestinal Pathogen

As malnutrition impairs immune functions in *Drosophila* (Vijendravarma et al., 2015), we asked if defined dietary modifications influence host responses to challenges with an intestinal pathogen. *Drosophila* is an established model for infection with the enteric pathogen *Vibrio cholerae* (Blow et al., 2005). To determine if diet altered survival time during a *V. cholerae* infection, we raised adult female flies on defined diets for ten days and measured survival after delivering a lethal infectious dose of *Vibrio.* We chose female flies for these studies, as they outlive male flies challenged with the same pathogen, allowing us to explore the effect of diet on survival to a greater extent. We found that flies on a holidic diet had a median survival of 49 hours after infection (Figure 3A and B). Supplementation with casein led to a slight decrease in median survival, while the other dietary modifications all showed an increase in median lifespan. Flies that were raised on increased glucose showed the most significant extension in survival during infection (□□=19.82, p<0.0001). These data establish that defined nutritional regimes influence the ability of *Drosophila* to combat an enteric infection, and in particular, that increased levels of glucose significantly elevate the survival times for adult flies.

**Figure 3:**
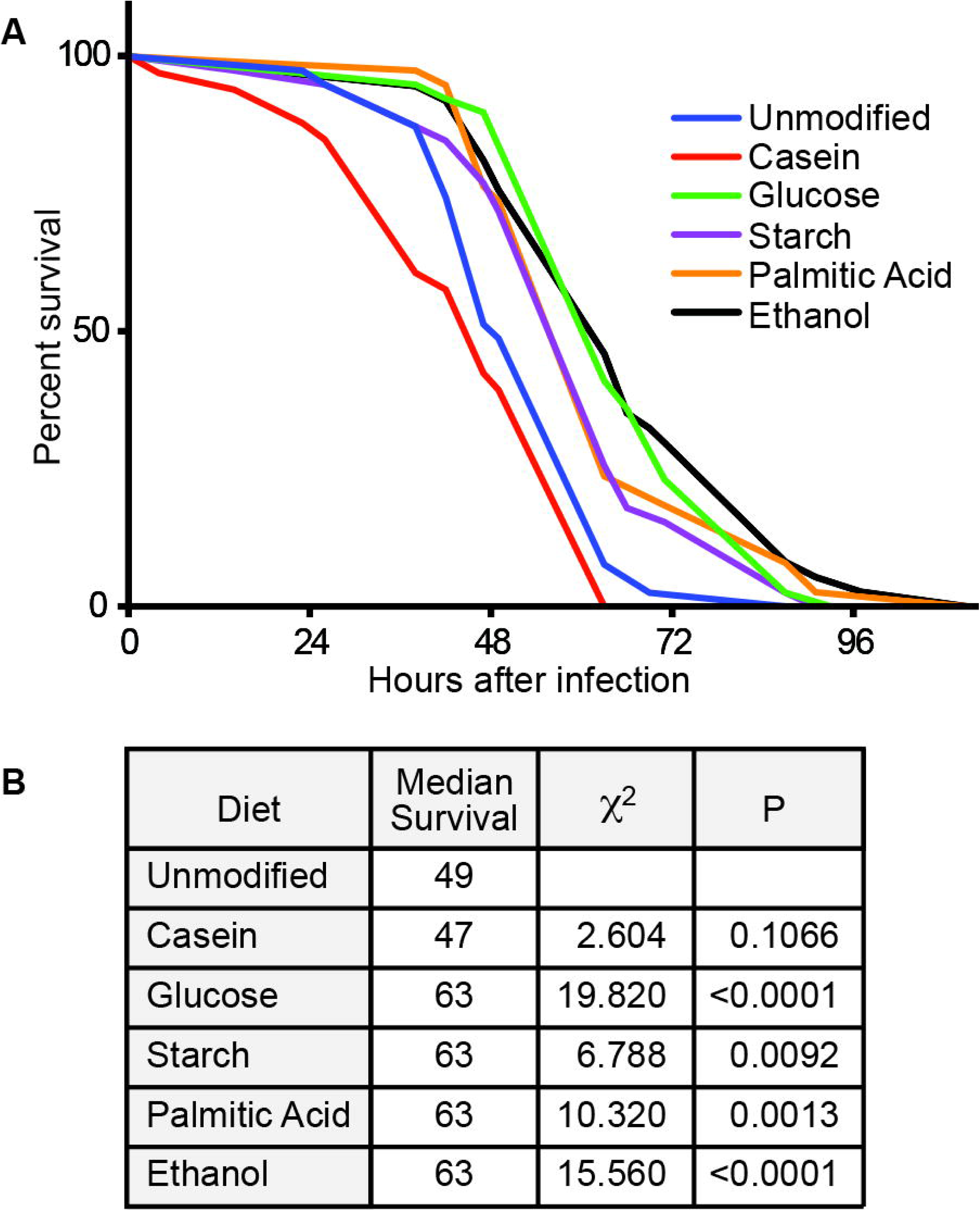
(A) Survival curves of female flies raised on unmodified or modified holidic diets for ten days and then challenged by a chronic infection with *V. cholerae* C6706. (B) Results of Logrank (Mantel-Cox) test of data in panel A. All □^2^ and p values are relative to wild type.

### Diet Influences Locomotion

As health and lifespan are commonly connected to physical activity, we asked if defined diets influence activity in adult *Drosophila.* For these assays, we compared locomotion in flies raised on a holidic diet to flies raised on the same diet supplemented with either glucose or ethanol. We chose glucose and ethanol, as both treatments extended median survival rates in adults. In each case, we trained flies with twelve-hour cycles of light and dark for five days, followed by five days of constant darkness. This approach allowed us to determine the effects of diet on activity, as well as the establishment and maintenance of circadian rhythms. We found that fly locomotion quickly adapted to defined cycles of light and dark irrespective of the diet, with peak activity levels after transitions to periods of light and sharp drops in locomotion after transitions to periods of dark (Figure 4A, days 1-5). For all treatments, adult flies maintained this behavioural pattern during the subsequent five days of constant darkness (Figure 4A, days 6-10). We observed a clear peak of activity that corresponded to a twenty-four hour period for days 1-5 and days 6-10, with a less prominent period of eight hours during days 1-5 irrespective of the diet (Figure 4B). Combined, these data show that the individual diets do not affect the ability of adult flies to maintain a circadian rhythm. However, examination of the data in Figures 4A and B suggest that diet influences general locomotion. To quantify the extent of this effect, we tallied total daily movements for flies raised on the respective diets. We found that supplementation with glucose greatly enhanced fly activity, while supplementation with ethanol had a sedative effect. Addition of extra glucose to the medium boosted daily activity levels by 39% relative to the control population on a holidic diet, while provision of moderate amounts of ethanol decreased daily locomotion by roughly 27% (Figure 4C).

**Figure 4:**
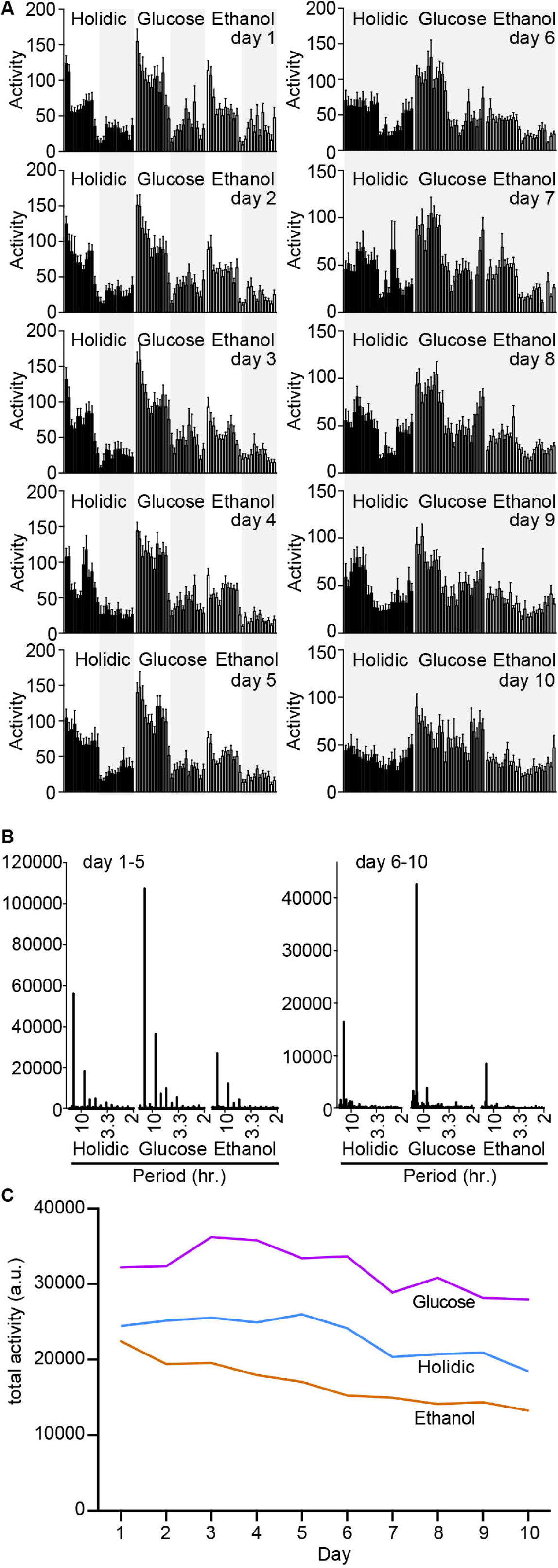
(A) Actograms showing average hourly activity of flies fed an unmodified diet (holidic), or a holidic diet supplemented with glucose or ethanol as indicated. Results are the mean of 20 individual measurements, and error bars indicate standard errors. Shaded areas indicate periods of darkness. (B) Periodograms of days 1-5 and 6-10 for flies raised on the respective diets. (C) Total daily activity of flies raised on the respective diets.

### Dietary Glucose Supplementation Alters Microbiota Composition and Increases Diversity

The studies detailed above uncover a number of effects of defined dietary modifications on the health of adult flies. As the microbiota of the host is known to affect these factors, we assessed the impact of defined diets on the intestinal microbiota. For these assays, we raised adults on modified diets and performed 16S DNA sequencing on bacterial DNA isolated from their intestinal tracts. Males and females raised on an unmodified holidic diet had similar microbiota that were dominated by the *Acetobacter* genus (Figure 5A). We found that simple alterations to this holidic diet resulted in profound changes in microbiota composition and diversity (Figure 5B). For example, when flies were raised on a diet supplemented with casein, the microbiota shifted to predominantly *Lactobacillus* species. In contrast, supplementation with glucose resulted in the largest increase in microbiota diversity (Shannon: Females=2.387, Males=1.789). We also noticed a different response between males and females to the same dietary modification, as seen for a diet supplemented with ethanol (Shannon: Females=2.383, Males=0.193). Our data suggest that both host diet and sex markedly impact the composition of intestinal microbiota, with supplementary glucose contributing to the greatest increase in species diversity.

**Figure 5:**
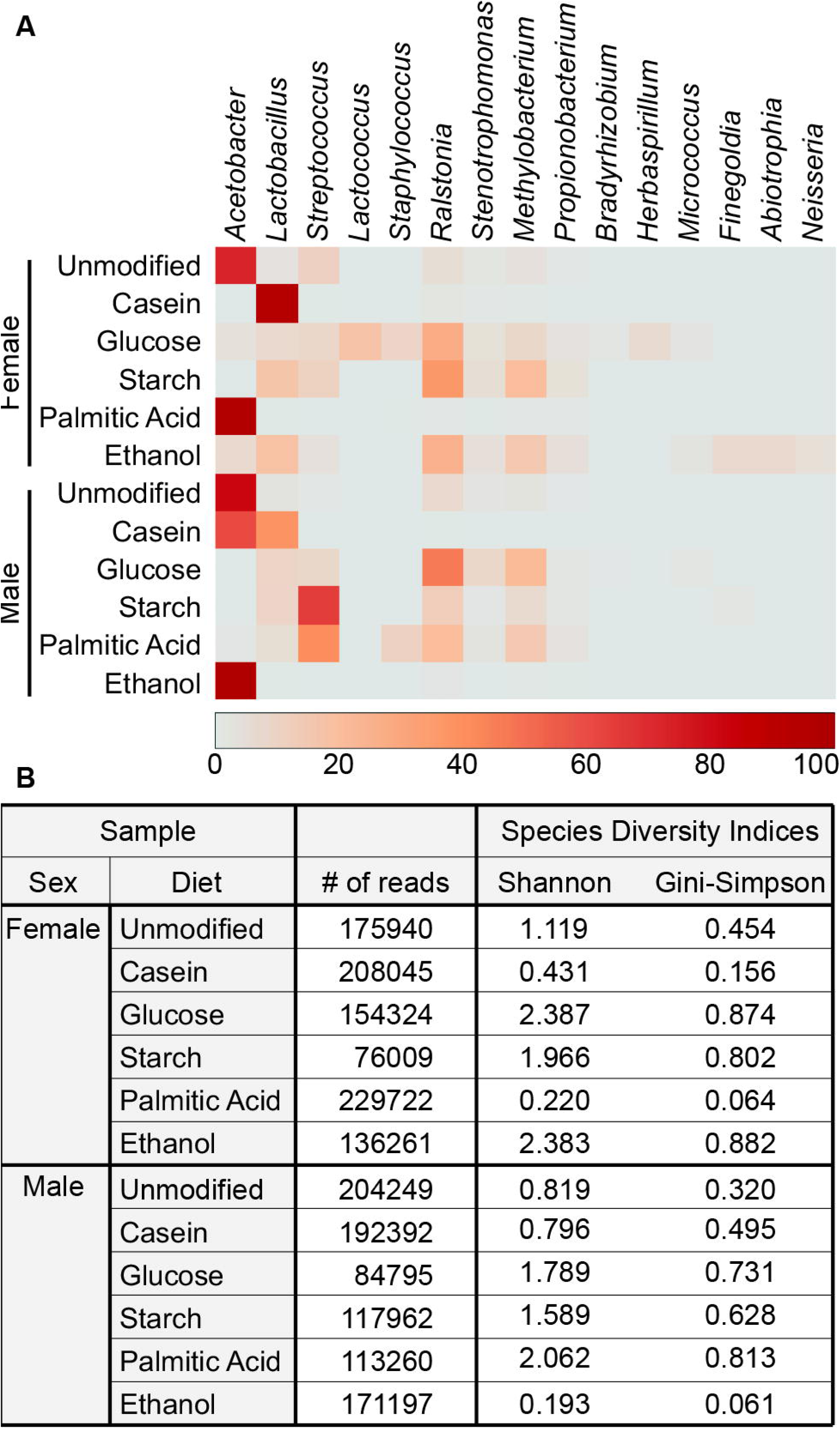
(A) Heat map summary showing abundance of bacterial genera present at greater than 1% in midguts of male and female flies raised on different diets for 10 days. Each sample consists of 5 flies. Abundance of each bacterial genus in a sample ranges from 0% (grey) to 100% (dark red) as indicated by the scale. (B) Summary showing the number of reads from 16S sequencing and the results from both Shannon and Gini-Simpson diversity values of each sample.

### Glucose Increases Lifespan and Survival to Infection Independent of the Microbiota

At this stage, our data reveal wide-ranging impacts of dietary glucose supplementation on adult flies, with significant effects on longevity, locomotion, energy stores, microfloral composition, and immunity. Given the established links between intestinal microflora diversity and host health (Norman et al., 2015), we asked if the microbiota is required for the beneficial effects of glucose supplementation on longevity. For these assays, we fed adult male flies an unmodified holidic diet or one supplemented with glucose and raised the flies under conventional or germ-free conditions. Consistent with recent reports (Clark et al., 2015; Petkau et al., 2014), we found that flies raised under germ-free conditions outlived their conventionally-reared counterparts (Figure 6A and B). Similar to our earlier experiments, we found that elevated dietary glucose increased the median lifespan of adult flies by 25% compared to an unmodified diet. Strikingly, we found that elimination of the microbiome did not affect the lifespan of flies raised on diets with elevated glucose, suggesting that glucose levels influence host longevity independently of the microbiome.

**Figure 6:**
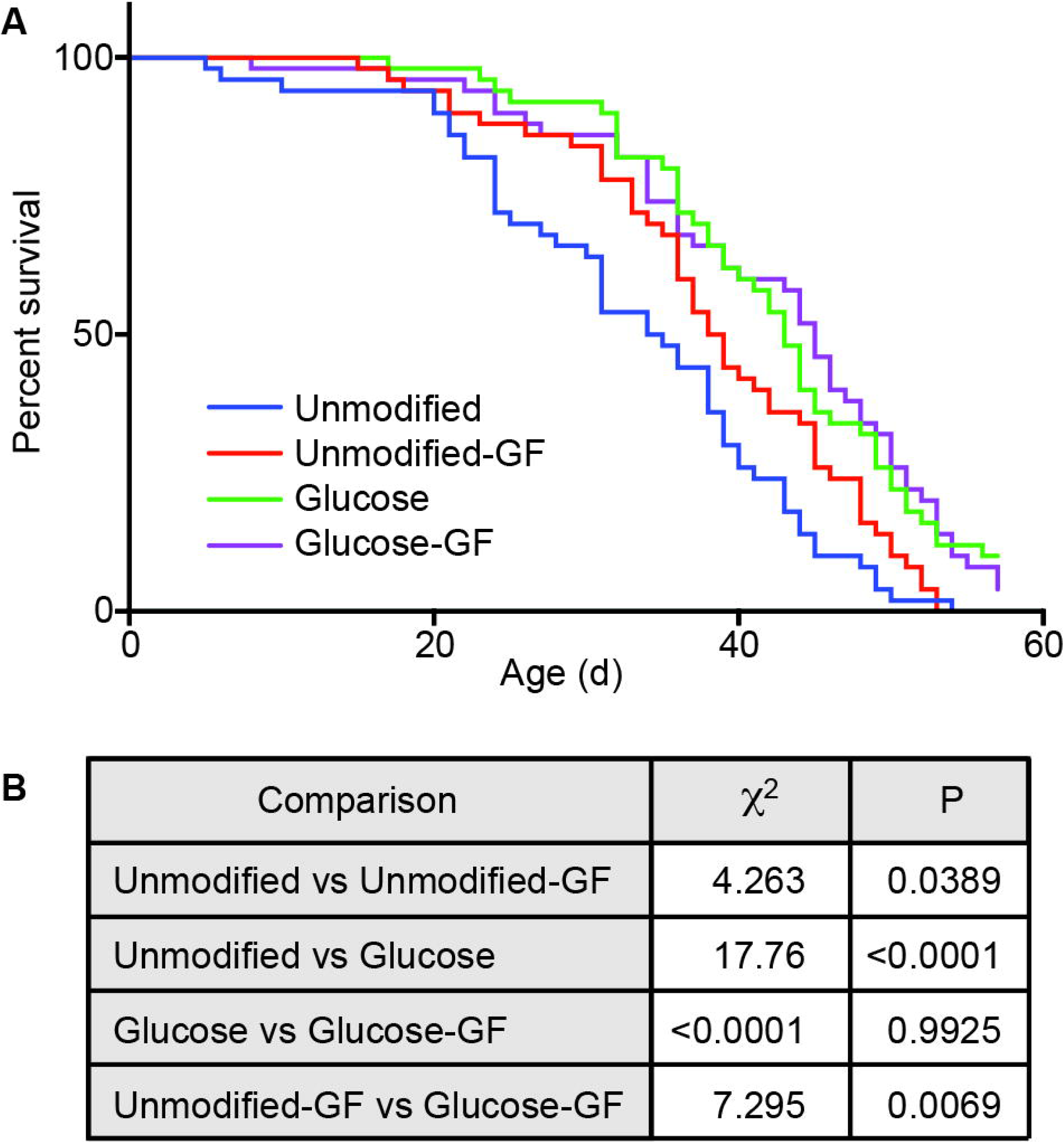
(A) Survival curves of males fed a holidic diet with or without glucose and raised under conventional or germ-free (GF) conditions. (B) Results of Log-rank (Mantel-Cox) test of data in panel A.

These unexpected observations prompted us to ask if the glucose-mediated improvements to survival after challenges with *Vibrio* challenges require a microbiome. To assess this, we fed adult *Drosophila* an unmodified holidic diet or one supplemented with glucose and raised the flies under conventional or germ-free conditions. We then measured survival after delivering a lethal infectious dose of *Vibrio*. As expected, we found that conventional flies on an unmodified diet had a median survival of 49.5 hours (Figure 7A). Removal of the microbiome significantly improved survival after infection with *Vibrio* (Figure 7B). As before, we found that elevated dietary glucose significantly improved survival compared to an unmodified diet (□□=17.390, p=<0.0001). Remarkably, elimination of the microflora did not alter the survival rates of flies raised on a high glucose diet and challenged with *Vibrio*. Combined, the data in Figures 6 and 7 establish that the microfloral shifts associated with transition to a high glucose diet are not essential for the immunological and lifespan benefits of such a diet.

**Figure 7:**
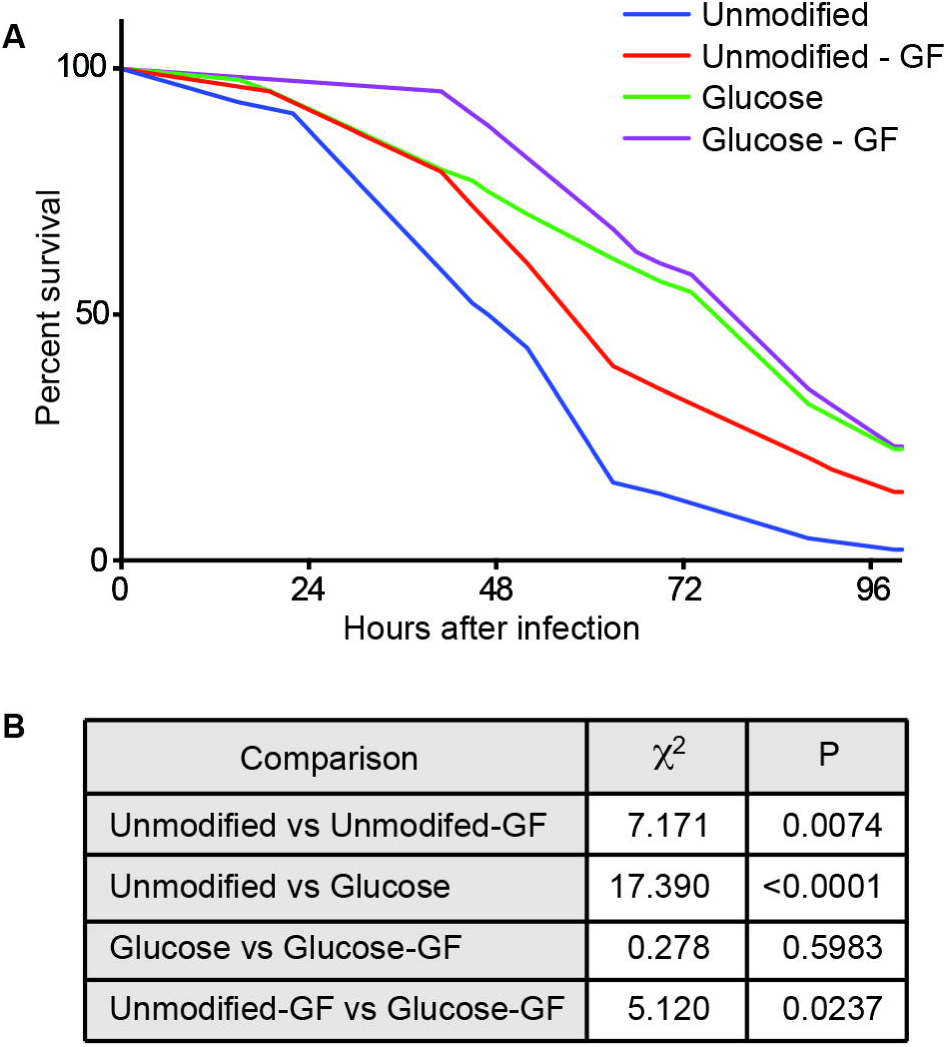
(A) Survival curves of chronically infected female flies fed a holidic diet with or without glucose and raised under conventional or germ-free (GF) conditions. (B) Results of Log-rank (Mantel-Cox) test of data in panel A.

### Increased Lifespan From Glucose is Independent of Intestinal Insulin Signaling

Our observation that glucose promotes antibacterial defenses independently of an intact intestinal microbiota suggests direct effects of glucose on host intestinal physiology. As previous studies showed that high-sugar diets cause insulin resistance in *Drosophila* (Musselman et al., 2011), and inhibition of insulin signalling in the gut promotes longevity (Biteau et al., 2010), we reasoned that elevated dietary glucose leads to insulin insensitivity in the intestine of adult flies, thereby extending the lifespans of the fly. To test this hypothesis, we generated a temperature sensitive *esgGAL4, GAL80^ts^/+*; *UASInR/+ (esg^ts^>InR) Drosophila* strain to control insulin receptor activity in midgut progentiors. In this strain, the combination of *esgGAL4* and *GAL80^ts^* transgenic elements induce insulin receptor (InR) activity in midgut progenitors of adult flies at the restrictive temperature of 29°C. As described in a previous study (Biteau et al., 2010), activation of the insulin receptor decreased the lifespans of adult flies compared to *esgGAL4, GAL80^ts^/+ (esg^ts^/+)* (Figure 8A and B). However, we found that *esg^ts^>InR* flies raised on a diet with added glucose significantly outlived *esg^ts^>InR* counterparts raised on an unmodified diet. In fact, the lifespan extensions observed upon addition of glucose were comparable for *esg^ts^>InR* and *esg^ts^* controls (Figure 8B). These data suggest that a high glucose diet extends adult *Drosophila* lifespan independent of insulin receptor activity in the gut.

**Figure 8:**
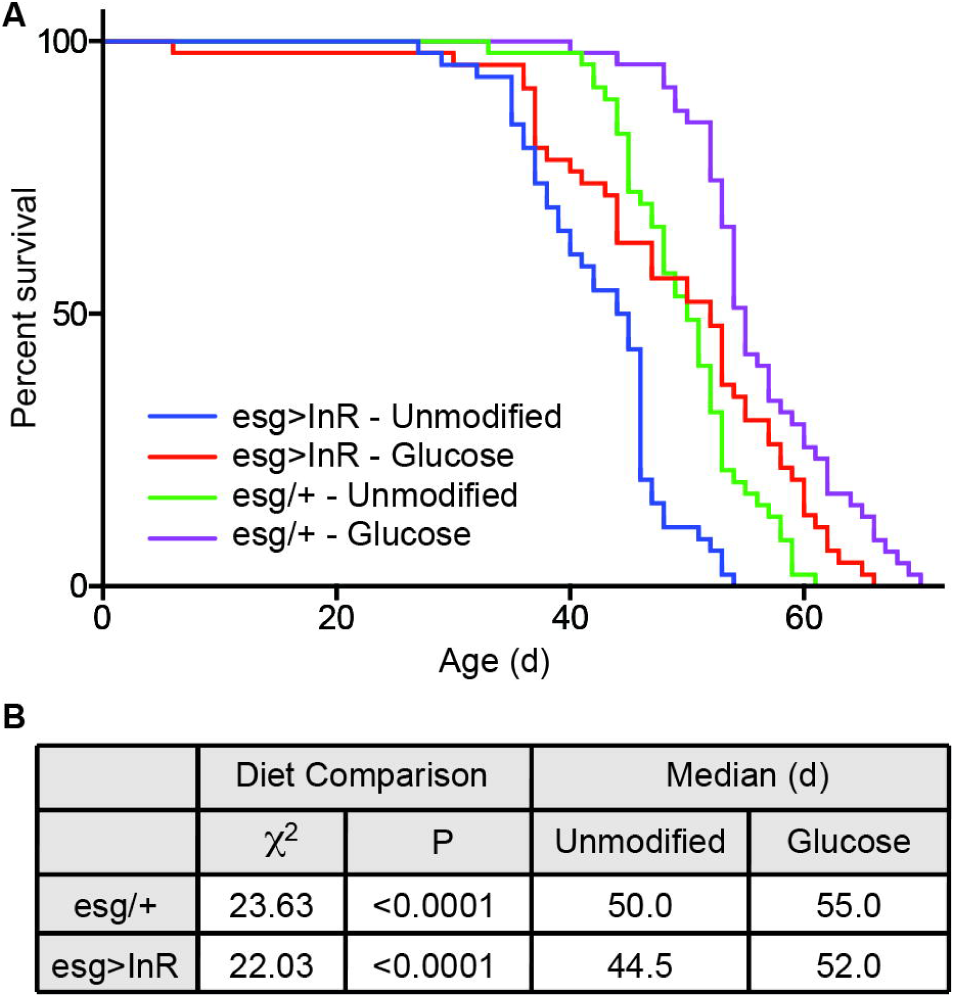
(A) Survival curves of male *esg>InR* or *esg/+* flies raised on an unmodified holidic diet or one supplemented with glucose. (B) Results of Log-rank (Mantel-Cox) test of data in panel A.

## DISCUSSION

The acquisition and allocation of nutrients is essential for maintenance of the intricate cellular processes that define multicellular life. Nutritional imbalances are at the heart of global health challenges that range from malnutrition to diabetes and meaningful solutions require a complete appreciation of the effects of complex diets on host wellbeing. We used *Drosophila melanogaster* as a tool to model the longitudinal consequences of defined diets in a common experimental system. The holidic diet allowed us to manipulate relative nutrient levels with far more precision than has been possible previously. Apart from the potentially whimsical observation of a proverbial “sugar rush”, we noticed a number of benefits of elevated levels of dietary glucose for adult flies. Increases to the levels of dietary glucose extended longevity, improved resistance to enteric infection, and increased diversity of the intestinal microbiota. The range of phenotypes suggests broad physiological responses to altered levels of dietary glucose.

Consistent with two recent reports (Clark et al., 2015; Petkau et al., 2014), we found that elimination of the adult microflora extended the lifespan of flies raised on a holidic diet. However, it is important to emphasize that the relationship between microfloral presence and fly longevity is by no means resolved, as other groups reported negative (Brummel et al., 2004) or neutral (Ren et al., 2007) effects of germ-free culture conditions on adult longevity. We consider it possible that the phenotypic variation between labs reflects differences between the respective *Drosophila* genotypes, diets, or microbiomes. Despite the phenotypic similarities between flies raised under germ-free conditions and flies raised on a high glucose diet, we established that glucose does not require a microbiome to improve immunity or longevity. Instead, our data suggest that host-intrinsic responses are key to the benefits of elevated levels of dietary glucose.

At first glance, the health benefits of elevated glucose appear incongruous with conventional dietary wisdoms, as chronic ingestion of carbohydrates is linked to metabolic disorders such as diabetes, and caloric restriction enhances longevity in many model organisms. However, recent studies established that dietary ratios of protein and carbohydrate (P:C) exert a substantial influence on the health of flies and mice, with several phenotypic benefits for animals raised on diets with low P:C ratios (Bruce et al., 2013; Grandison et al., 2009; Lee et al., 2008; Mirzaei et al., 2014; Piper et al., 2011; Solon-Biet et al., 2014; Solon-Biet et al., 2015a; Solon-Biet et al., 2015c). Consistent with this hypothesis, we showed that addition of glucose, an effective drop in P:C ratio, boosted longevity, while addition of casein, an effective increase in P:C ratio, reduced the lifespan of adult flies. Our studies extended these observations to show that low P:C ratios improve immune responses to an intestinal pathogen, and that elevated P:C ratios are detrimental to survival after infection with *Vibrio cholerae.* Of note, the strength of the phenotypes differed for male and female flies, suggesting endocrinological regulation of dietary effects on lifespan.

To date, there are few studies that explore interplays between nutrient access, microfloral composition and animal health. In a preliminary set of experiments, we found that germ-free flies resisted chronic intestinal challenges with *Vibrio cholerae* better than conventional flies. The fly response to enteric infection is remarkably sophisticated and includes the expression of bactericidal AMPs through the Immune Deficiency (IMD) response (Lemaitre and Miguel-Aliaga, 2013). A previous study demonstrated that mutations in the IMD pathway extend the lifespan of flies challenged with *Vibrio cholerae* (Wang et al., 2013), suggesting deleterious contributions from IMD to the pathogenesis of *Vibrio cholerae.* As intestinal IMD activity is lower in flies raised under germ-free conditions (Broderick et al., 2014), we speculate that germ-free flies are better equipped to deal with *Vibrio* infections due to diminished intestinal IMD activity. However, our data do not exclude alternative explanations such as effects of glucose on the expression of pathogenicity factors by *Vibrio*. Future studies are required to determine the mechanistic basis for dietary alterations of intestinal immune responses.

As nutritional status influences intestinal immunity in *Drosophila* (Vijendravarma et al., 2015), we asked if dietary modifications improve immune responses to *Vibrio* by modification of the intestinal microflora. Several studies demonstrated that diet influences composition of the fly’s microbiota (Staubach et al., 2013; Wong et al., 2013). However, these studies relied on substantial alterations to partially defined diets. We used the holidic diet to examine the effects of specific macronutrients on microbiome makeup. Our results showed that remarkably simple dietary changes, such as alterations to relative glucose or protein levels, drastically alter the microbiota of adult flies. However, at least in the case of glucose, these modifications do not directly influence the lifespan or immune response, as germ-free flies and their conventionally reared counterparts have indistinguishable viability profiles when raised on diets with elevated glucose.

As a cautionary note, we cannot exclude that some dietary modifications act at least partially through effects on the microbiota in *Drosophila.* For example, a recent study established that the commensal fungus *Isatchenkia orientalis* promotes amino acid uptake in nutritionally deprived *Drosophila* (Yamada et al., 2015). In our study, we found that flies fed a diet supplemented with moderate amounts of ethanol lived longer and survived infection better than control flies raised on an unmodified diet. Since wild flies develop in and consume decomposed fruit, ethanol is likely a common constituent of their environment. As ethanol is a fuel source for *Acetobacter*, a prominent fly commensal that modifies insulin and TOR signals in the midgut (Shin et al., 2011; Storelli et al., 2011), it remains possible that the lifespan extension we see from ethanol depends on the microbiota.

Since we observed benefits from glucose independent of the microbiota, we were interested in the host response to glucose that extends lifespan. High dietary glucose lowers insulin sensitivity in flies (Musselman et al., 2011), and decreased insulin activity in the gut increases lifespan in the fly (Biteau et al., 2010). These observations led us to speculate that glucose increases longevity through reduced insulin receptor activity in the gut. However, our results showed that overexpression of the insulin receptor in intestinal progenitor cells did not impair the lifespan benefits of added glucose, suggesting that outputs from the gut insulin receptor do not influence the benefits of elevated dietary glucose. Animals respond to their nutritional environment through complex signal transduction pathways such as TOR and insulin in several organs, and both pathways influence lifespan and health in numerous model organisms (Harrison et al., 2009; Kapahi and Zid, 2004; Kapahi et al., 2004; Scialo et al., 2015; Tatar et al., 2003; Vellai et al., 2003). In the case of flies, the gut, fat body, and insulin-producing neurons coordinate the uptake and distribution of macronutrients (Buch et al., 2008). A more extensive analysis of the physiological benefits of increased glucose will likely require dissection of TOR and insulin responses in these tissues.

In summary, this study complements an emerging body of literature that low dietary P:C ratios extend the lifespan of *Drosophila.* In addition, we show that alterations to P:C ratios generate significant phenotypes in locomotion, immunity, and microfloral diversity. Despite the links between the intestinal microflora and animal health, we established that glucose acts directly on the host to increase lifespan and responses to *cholerae* infections. As physiological responses to diet are extensively conserved throughout the animal kingdom, we believe our findings may be of relevance to a general appreciation of the relationship between glucose consumption and animal health.

## MATERIALS AND METHODS

### Fly Husbandry

All experiments were performed with virgin female and male adult flies raised at 29°C. *w^1118^* flies were used as a control genotype unless otherwise mentioned. The *esgGAL4* flies described in this study have been described elsewhere (Buchon et al., 2009). The wild *Drosophila melanogaster* population was derived from a single pregnant female captured in EF’s kitchen in Edmonton (Canadan) in the summer of 2014. Adult flies were raised on a holidic medium developed by Piper *et al.* using the Oaa stock recipe and 100 mM biologically available nitrogen (Piper et al., 2014). Modifications to this diet include: the additions of simple sugar (100 g/L *D*-glucose), complex sugar (50 g/L starch), protein (70 g/L casein), fatty acid (50 g/L palmitic acid), and alcohol (1% ethanol). Germ-free flies were generated by raising adults on food supplemented with an antibiotic cocktail (100 μg/mL ampicillin, 50 μg/mL vancomycin, 100 μg/mL neomycin, and 100 μg/mL metronidazole). Survival assays were performed with approx. 50 flies housed at 10 flies/vial and transferred to fresh vials weekly.

### Metabolic Assays

All metabolic assays were performed as described in (Wong et al., 2014) in 96-well plates using commercial kits: the DC Protein Assay kit (Bio-Rad, 500-0116), Triglyceride Assay kit (Sigma, TG-5-RB), and Glucose (GO) Assay kit (Sigma, GAGO20). Colorimetric readings were obtained using a microplate spectrophotometer (PerkinElmer, Envision Multilabel Reader).

### Locomotion Assay

Holidic medium was added to small glass tubes capped with a plastic stopper. A single fly was placed in each tube. The tube was then closed with yarn and then placed into monitors of the TriKinetics DAMSystem. Fly activity was monitored for 10 days at 23°C. A 12 h light/ 12 h dark cycle for 5 days was used for circadian training followed by 5 days in permanent dark.

### Infection Protocol

Samples of 40-50 adult female flies at 15 flies/vial, were raised for 10 days at 29°C. Flies were starved by placing in empty vials for 2 h prior to infection. Flies were then maintained in vials containing a cellulose acetate plug infused with 3 mL of *V. cholerae* C6706 (10^8^-10^9^ cells/mL) in LB, and viability was monitored over a 5-day period.

### Microbiota sequencing

Samples of 10 adult male or female flies were raised for 10 days at 29°C. Intestinal tracts were dissected as described elsewhere and bacterial genomic DNA was isolated with the Ultraclean Microbial DNA Isolation Kit (MO-BIO, 12224). Bacterial 16S DNA was amplified with primers AGAGTTTGATCCTGGCTCAG (forward) and GGCTACCTTGTTACGACTT (reverse). Samples were purified with a QIAquick PCR Purification Kit (Qiagen, 28104). Samples were prepared for sequencing using a Nextera XT DNA Library Preparation Kit (Illumina, FC-131-1024), and DNA libraries were sequenced using a MiSeq Desktop Sequencer (Illumina). Taxonomy assignment was based on the SILVA SSU Ref NR99 database release 115, using in-house developed software.

## ACKNOWLEDGEMENTS

Jeff Reeve (Alberta Transplant Applied Genomics Centre) assisted with preparation of actograms. The Applied Genomics Core at the University of Alberta assisted with microbial sequencing. *esgGAL4, GAL80^ts^* flies were provided by Bruno Lemaitre, and *UAS-InR* flies were provided by Kirst King-Jones.

## REFERENCE

Biteau, B., Karpac, J., Supoyo, S., Degennaro, M., Lehmann, R., Jasper, H., 2010. Lifespan extension by preserving proliferative homeostasis in Drosophila. PLoS Genet 6, el001159.

Blow, N.S., Salomon, R.N., Garrity, K., Reveillaud, I., Kopin, A., Jackson, F.R., Watnick, P.I., 2005. Vibrio cholerae infection of Drosophila melanogaster mimics the human disease cholera. PLoS Pathog 1, e8.

Broderick, N.A., Buchon, N., Lemaitre, B., 2014. Microbiota-induced changes in drosophila melanogaster host gene expression and gut morphology. MBio 5, e01117–01114.

Broderick, N.A., Lemaitre, B., 2012. Gut-associated microbes of Drosophila melanogaster. Gut Microbes 3, 307–321.

Bruce, K.D., Hoxha, S., Carvalho, G.B., Yamada, R., Wang, H.D., Karayan, P., He, S., Brummel, T., Kapahi, P., Ja, W.W., 2013. High carbohydrate-low protein consumption maximizes Drosophila lifespan. Exp Gerontol 48,1129–1135.

Brummel, T., Ching, A., Seroude, L., Simon, A.F., Benzer, S., 2004. Drosophila lifespan enhancement by exogenous bacteria. Proc Natl Acad Sei U S A 101,12974–12979.

Buch, S., Melcher, C., Bauer, M., Katzenberger, J., Pankratz, M.J., 2008. Opposing effects of dietary protein and sugar regulate a transcriptional target of Drosophila insulin-like peptide signaling. Cell Metab 7, 321–332.

Buchon, N., Broderick, N.A., Chakrabarti, S., Lemaitre, B., 2009. Invasive and indigenous microbiota impact intestinal stem cell activity through multiple pathways in Drosophila. Genes Dev 23, 2333–2344.

Buchon, N., Broderick, N.A., Lemaitre, B., 2013. Gut homeostasis in a microbial world: insights from Drosophila melanogaster. Nat Rev Microbiol 11, 615–626.

Clark, R.I., Salazar, A., Yamada, R., Fitz-Gibbon, S., Morselli, M., Alcaraz, J., Rana, A., Rera, M., Pellegrini, M., Ja, W.W., Walker, D.W., 2015. Distinct Shifts in Microbiota Composition during Drosophila Aging Impair Intestinal Function and Drive Mortality. Cell Rep 12,1656–1667.

Colman, R.J., Beasley, T.M., Kemnitz, J.W., Johnson, S.C., Weindruch, R., Anderson, R.M., 2014. Caloric restriction reduces age-related and all-cause mortality in rhesus monkeys. Nat Commun 5, 3557.

Dilova, I., Easlon, E., Lin, S.J., 2007. Calorie restriction and the nutrient sensing signaling pathways. Cell Mol Life Sci 64, 752–767.

Erkosar, B., Defaye, A., Bozonnet, N., Puthier, D., Royet, J., Leulier, F., 2014. Drosophila microbiota modulates host metabolic gene expression via IMD/NF-kappaB signaling. PLoS One 9, e94729.

Erkosar, B., Leulier, F., 2014. Transient adult microbiota, gut homeostasis and longevity: novel insights from the Drosophila model. FEBS Lett 588, 4250–4257.

Flint, H.J., Scott, K.P., Louis, P., Duncan, S.H., 2012. The role of the gut microbiota in nutrition and health. Nat Rev Gastroenterol Hepatol 9, 577–589.

Fontana, L., Partridge, L., 2015. Promoting health and longevity through diet: from model organisms to humans. Cell 161,106–118.

Fontana, L., Partridge, L., Longo, V.D., 2010. Extending healthy life span-from yeast to humans. Science 328, 321–326.

Grandison, R.C., Piper, M.D., Partridge, L., 2009. Amino-acid imbalance explains extension of lifespan by dietary restriction in Drosophila. Nature 462, 1061–1064.

Harrison, D.E., Strong, R., Sharp, Z.D., Nelson, J.F., Astle, C.M., Flurkey, K., Nadon, N.L., Wilkinson, J.E., Frenkel, K., Carter, C.S., Pahor, M., Javors, M.A., Fernandez, E., Miller, R.A., 2009. Rapamycin fed late in life extends lifespan in genetically heterogeneous mice. Nature 460,392–395.

Harvie, M.N., Pegington, M., Mattson, M.P., Frystyk, J., Dillon, B., Evans, G., Cuzick, J., Jebb, S.A., Martin, B., Cutler, R.G., Son, T.G., Maudsley, S., Carlson, O.D., Egan, J.M., Flyvbjerg, A., Howell, A., 2011. The effects of intermittent or continuous energy restriction on weight loss and metabolic disease risk markers: a randomized trial in young overweight women. Int J Obes (Lond) 35, 714–727.

Honjoh, S., Yamamoto, T., Uno, M., Nishida, E., 2009. Signalling through RHEB-1 mediates intermittent fasting-induced longevity in C. elegans. Nature 457, 726–730.

Hooper, L.V., Littman, D.R., Macpherson, A.J., 2012. Interactions between the microbiota and the immune system. Science 336,1268–1273.

Jakubowicz, D., Barnea, M., Wainstein, J., Froy, O., 2013. Effects of caloric intake timing on insulin resistance and hyperandrogenism in lean women with polycystic ovary syndrome. Clin Sci (Lond) 125, 423–432.

Kapahi, P., Zid, B., 2004. TOR pathway: linking nutrient sensing to life span. Sci Aging Knowledge Environ 2004, PE34.

Kapahi, P., Zid, B.M., Harper, T., Koslover, D., Sapin, V., Benzer, S., 2004. Regulation of lifespan in Drosophila by modulation of genes in the TOR signaling pathway. Curr Biol 14, 885–890.

Lee, D., Hwang, W., Artan, M., Jeong, D.E., Lee, S.J., 2015. Effects of nutritional components on aging. Aging Cell 14, 8–16.

Lee, K.P., Simpson, S.J., Clissold, F.J., Brooks, R., Ballard, J.W., Taylor, P.W., Soran, N., Raubenheimer, D., 2008. Lifespan and reproduction in Drosophila: New insights from nutritional geometry. Proc Natl Acad Sci U S A 105, 2498–2503.

Lemaitre, B., Miguel-Aliaga, I., 2013. The digestive tract of Drosophila melanogaster. Annu Rev Genet 47, 377–404.

Ma, D., Storelli, G., Mitchell, M., Leulier, F., 2015. Studying host-microbiota mutualism in Drosophila: Harnessing the power of gnotobiotic flies. Biomed J 38, 285–293.

Mair, W., Goymer, P., Pletcher, S.D., Partridge, L., 2003. Demography of dietary restriction and death in Drosophila. Science 301,1731–1733.

Mattison, J.A., Roth, G.S., Beasley, T.M., Tilmont, E.M., Handy, A.M., Herbert, R.L., Longo, D.L., Allison, D.B., Young, J.E., Bryant, M., Barnard, D., Ward, W.F., Qi, W., Ingram, D.K., de Cabo, R., 2012. Impact of caloric restriction on health and survival in rhesus monkeys from the NIA study. Nature 489, 318–321.

Mattson, M.P., Allison, D.B., Fontana, L., Harvie, M., Longo, V.D., Malaisse, W.J., Mosley, M., Notterpek, L., Ravussin, E., Scheer, F.A., Seyfried, T.N., Varady, K.A., Panda, S., 2014. Meal frequency and timing in health and disease. Proc Natl Acad Sci U S A 111, 16647–16653.

Miller, R.A., Buehner, G., Chang, Y., Harper, J.M., Sigler, R., Smith-Wheelock, M., 2005. Methionine-deficient diet extends mouse lifespan, slows immune and lens aging, alters glucose, T4, IGF-I and insulin levels, and increases hepatocyte MIF levels and stress resistance. Aging Cell 4,119–125.

Mirzaei, H., Suarez, J.A., Longo, V.D., 2014. Protein and amino acid restriction, aging and disease: from yeast to humans. Trends Endocrinol Metab 25, 558–566.

Musselman, L.P., Fink, J.L., Narzinski, K., Ramachandran, P.V., Hathiramani, S.S., Cagan, R.L., Baranski, T.J., 2011. A high-sugar diet produces obesity and insulin resistance in wild-type Drosophila. Dis Model Mech 4, 842–849.

Nakagawa, S., Lagisz, M., Hector, K.L., Spencer, H.G., 2012. Comparative and meta-analytic insights into life extension via dietary restriction. Aging Cell 11, 401–409.

Newell, P.D., Douglas, A.E., 2014. Interspecies interactions determine the impact of the gut microbiota on nutrient allocation in Drosophila melanogaster. Appl Environ Microbiol 80, 788–796.

Norman, J.M., Handley, S.A., Baldridge, M.T., Droit, L., Liu, C.Y., Keller, B.C., Kambal, A., Monaco, C.L., Zhao, G., Fleshner, P., Stappenbeck, T.S., McGovern, D.P., Keshavarzian, A., Mutlu, E.A., Sauk, J., Gevers, D., Xavier, R.J., Wang, D., Parkes, M., Virgin, H.W., 2015. Disease-specific alterations in the enteric virome in inflammatory bowel disease. Cell 160, 447–460.

Osawa, R., Blanshard, W.H., O'Callaghan, P.G., 1992. Microflora of the pouch of the koala (Phascolarctos cinereus). J Wildl Dis 28, 276–280.

Petkau, K., Parsons, B.D., Duggal, A., Foley, E., 2014. A deregulated intestinal cell cycle program disrupts tissue homeostasis without affecting longevity in Drosophila. J Biol Chem 289,28719–28729.

Piper, M.D., Blanc, E., Leitao-Goncalves, R., Yang, M., He, X., Linford, N.J., Hoddinott, M.P., Hopfen, C., Soultoukis, G.A., Niemeyer, C., Kerr, F., Pletcher, S.D., Ribeiro, C., Partridge, L., 2014. A holidic medium for Drosophila melanogaster. Nat Methods 11,100–105.

Piper, M.D., Partridge, L., Raubenheimer, D., Simpson, S.J., 2011. Dietary restriction and aging: a unifying perspective. Cell Metab 14,154–160.

Ren, C., Webster, P., Finkel, S.E., Tower, J., 2007. Increased internal and external bacterial load during Drosophila aging without life-span trade-off. Cell Metab 6,144–152.

Round, J.L., Mazmanian, S.K., 2009. The gut microbiota shapes intestinal immune responses during health and disease. Nat Rev Immunol 9, 313–323.

Scialo, F., Sriram, A., Naudi, A., Ayala, V., Jove, M., Pamplona, R., Sanz, A., 2015. Target of rapamycin activation predicts lifespan in fruit flies. Cell Cycle, 0.

Sharon, G., Segal, D., Ringo, J.M., Hefetz, A., Zilber-Rosenberg, I., Rosenberg, E., 2010. Commensal bacteria play a role in mating preference of Drosophila melanogaster. Proc Natl Acad Sci U S A 107, 20051–20056.

Shin, S.C., Kim, S.H., You, H., Kim, B„ Kim, A.C., Lee, K.A., Yoon, J.H., Ryu, J.H., Lee, W.J., 2011. Drosophila microbiome modulates host developmental and metabolic homeostasis via insulin signaling. Science 334, 670–674.

Simpson, S.J., Le Couteur, D.G., Raubenheimer, D., 2015. Putting the balance back in diet. Cell 161,18–23.

Simpson, S.J., Raubenheimer, D., 2009. Macronutrient balance and lifespan. Aging (Albany NY) 1, 875–880.

Solon-Biet, S.M., McMahon, A.C., Ballard, J.W., Ruohonen, K., Wu, L.E., Cogger, V.C., Warren, A., Huang, X., Pichaud, N., Melvin, R.G., Gokarn, R., Khalil, M., Turner, N., Cooney, G.J., Sinclair, D.A., Raubenheimer, D., Le Couteur, D.G., Simpson, S.J., 2014. The ratio of macronutrients, not caloric intake, dictates cardiometabolic health, aging, and longevity in ad libitum-fed mice. Cell Metab 19,418–430.

Solon-Biet, S.M., Mitchell, S.J., Coogan, S.C., Cogger, V.C., Gokarn, R., McMahon, A.C., Raubenheimer, D., de Cabo, R., Simpson, S.J., Le Couteur, D.G., 2015a. Dietary Protein to Carbohydrate Ratio and Caloric Restriction: Comparing Metabolic Outcomes in Mice. Cell Rep 11, 1529–1534.

Solon-Biet, S.M., Mitchell, S.J., de Cabo, R., Raubenheimer, D., Le Couteur, D.G., Simpson, S.J., 2015b. Macronutrients and caloric intake in health and longevity. J Endocrinol 226, R17–28.

Solon-Biet, S.M., Walters, K.A., Simanainen, U.K., McMahon, A.C., Ruohonen, K., Ballard, J.W., Raubenheimer, D., Handelsman, D.J., Le Couteur, D.G., Simpson, S.J., 2015c. Macronutrient balance, reproductive function, and lifespan in aging mice. Proc Natl Acad Sci U S A 112, 3481–3486.

Staubach, F., Baines, J.F., Kunzel, S., Bik, E.M., Petrov, D.A., 2013. Host species and environmental effects on bacterial communities associated with Drosophila in the laboratory and in the natural environment. PLoS One 8, e70749.

Storelli, G., Defaye, A., Erkosar, B., Hols, P., Royet, J., Leulier, F., 2011. Lactobacillus plantarum promotes Drosophila systemic growth by modulating hormonal signals through TOR-dependent nutrient sensing. Cell Metab 14, 403–414.

Tatar, M., Bartke, A., Antebi, A., 2003. The endocrine regulation of aging by insulin-like signals. Science 299,1346–1351.

Tatar, M., Post, S., Yu, K., 2014. Nutrient control of Drosophila longevity. Trends Endocrinol Metab 25, 509–517.

Vellai, T., Takacs-Vellai, K., Zhang, Y., Kovacs, A.L., Orosz, L., Muller, F., 2003. Genetics: influence of TOR kinase on lifespan in C. elegans. Nature 426, 620.

Vijendravarma, R.K., Narasimha, S., Chakrabarti, S., Babin, A., Kolly, S., Lemaitre, B., Kawecki, T.J., 2015. Gut physiology mediates a trade-off between adaptation to malnutrition and susceptibility to food-borne pathogens. Ecol Lett 18,1078–1086.

Wang, Z., Hang, S., Purdy, A.E., Watnick, P.I., 2013. Mutations in the IMD pathway and mustard counter Vibrio cholerae suppression of intestinal stem cell division in Drosophila. MBio 4, e00337–00313.

Wlodarska, M., Kostic, A.D., Xavier, R.J., 2015. An integrative view of microbiome-host interactions in inflammatory bowel diseases. Cell Host Microbe 17, 577–591.

Wong, A.C., Chaston, J.M., Douglas, A.E., 2013. The inconstant gut microbiota of Drosophila species revealed by 16S rRNA gene analysis. ISME J 7,1922–1932.

Wong, A.C., Dobson, A.J., Douglas, A.E., 2014. Gut microbiota dictates the metabolic response of Drosophila to diet. J Exp Biol 217,1894–1901.

Wu, Z., Song, L., Liu, S.Q., Huang, D., 2013. Independent and additive effects of glutamic acid and methionine on yeast longevity. PLoS One 8, e79319.

Yamada, R., Deshpande, S.A., Bruce, K.D., Mak, E.M., Ja, W.W., 2015. Microbes Promote Amino Acid Harvest to Rescue Undernutrition in Drosophila. Cell Rep.

